# Robust and Gradient Thickness Porous Membranes for *In Vitro* Modeling of Physiological Barriers

**DOI:** 10.1101/2020.05.07.083188

**Authors:** Shayan Gholizadeh, Zahra Allahyari, Robert Carter, Luis F. Delgadillo, Marine Blaquiere, Frederic Nouguier-Morin, Nicola Marchi, Thomas R. Gaborski

## Abstract

Porous membranes are fundamental elements for tissue-chip barrier and co-culture models. However, the exaggerated thickness of commonly available membranes impedes an accurate *in vitro* reproduction of the biological multi-cellular continuum as it occurs *in vivo*. Existing techniques to fabricate membranes such as solvent cast, spin-coating, sputtering and PE-CVD result in uniform thickness films. To understand critical separation distances for various barrier and co-culture models, a gradient thickness membrane is needed. Here, we developed a robust method to generate ultrathin porous parylene C (UPP) membranes not just with precise thicknesses down to 300 nm, but with variable gradients in thicknesses, while at the same time having porosities up to 25%. We also show surface etching and increased roughness lead to improved cell attachment. Next, we examined the mechanical properties of UPP membranes with varying porosity and thickness and fit our data to previously published models, which can help determine practical upper limits of porosity and lower limits of thickness. Lastly, we validate a straightforward approach allowing the successful integration of the UPP membranes into a prototyped 3D-printed scaffold enabling *in vitro* barrier modeling and investigation of cell-cell interplay over variable distances using thickness gradients.

## 1. Introduction

Modeling, *in vitro*, the structure of specific cell complexes and their environments is of utmost importance to examine the molecular mechanisms governing cell-to-cell interplay and to perform robust pharmacological studies.^[1–3]^ In particular, modeling physiological vascular barriers is of fundamental translational value, and novel solutions for *in vitro* study are continuously emerging with the goal of an effective integration to, or even to by-pass, *in vivo* approaches. When modeling vascular barriers, selecting the appropriate model cells is crucial as demonstrated by testing primary or immortalized endothelial cells and, more recently, human-derived pluripotent stem cells.^[4,5]^ However, this cellular-level advancement is not being matched with a research effort examining equally important aspects of *in vitro* modeling, specifically the development of porous membranes to mimic the nano-to-microscale level cell-to-cell interplay.^[6]^ Literature indicates that porous membranes are an essential component for compartmentalized cell co-cultures and to support a tissue barrier.^[6–15]^ However, most barrier models incorporate commercially available track-etched membranes such as those found in transwell, fiber-based, or micro-fluidic systems with a high thickness compared to the basal laminae *in vivo* (>10 µm vs. ~300 nm). By using available materials, the porosity cannot be adequately engineered, leading to low porosity, random pore distribution, and limited size control, potentially biasing physiological multi-cellular interplay, especially during co-culture conditions.^[16,17]^

The ideal membrane should allow necessary cell-cell communication and transport while providing the opportunity to modulate cell-substrate interactions through a change in porosity and pore geometry.^[18–20]^ In successful efforts to provide these properties, inorganic or polymeric ultrathin films have been developed and tested, providing physiologically relevant thickness, optical transparency, and controlled pore size.^[6,9,13,15,21–24]^ However, these solutions present significant fabrication or technical stumbling blocks that impede their integration into the *in vitro* modeling realm.

Limited active area for cell culture, time-consuming and costly process of backside etching of silicon wafers, and the use of toxic or corrosive solvents are among issues that need to be resolved. Importantly, existing techniques to fabricate membranes such as solvent casting, spin-coating, sputtering, and plasma-enhanced chemical vapor deposition (PE-CVD) result in uniform thickness films.^[25–28]^ In order to understand critical separation distances for various barrier and co-culture models, a gradient thickness membrane is needed. However, to the best of our knowledge, a continuous thickness gradient membrane that can be used for quantifying distance-dependent cell-cell communication in various tissue-on-a-chip and human barrier models have not been developed nor implemented.

Parylene C has been proposed as a material solution compatible with *in vitro* cell attachment and biological response, with a relatively low-cost fabrication over large surface areas with tunable thicknesses.^[29–35]^ Current protocols for parylene etching needs significant optimization, as they result in loss of pattern fidelity by changing membrane thickness.^[36–38]^ Therefore, precise pore geometry requires addition and subsequent removal of a hard mask which can be a time and resource expensive process.^[29,35]^ Parylene also tends to curl as a thin film as a result of thickness-dependent stress variation, which requires the addition of an undesirable and interfering chip or grid-based support.

In the present study, we developed facile methods to generate ultrathin porous parylene C (UPP) membranes not just of precise thicknesses down to 300 nm, but with variable gradients in thicknesses, all with relatively high porosities. We examined the mechanical properties of UPP membranes with varying porosity and thickness and fit our data to available models. We developed a straightforward approach to reinforce the ultrathin membranes while maintaining a large uninterrupted central region and integrated the UPP membrane into a prototyped 3D printed scaffold for tissue-chip devices.

## 2. Results and discussion

### 2.1. Membrane Fabrication and Scanning Electron Microscopy Characterizations

In order to maximize reproducibility, we sought to develop a simplified and reliable process for UPP membrane fabrication (Figure 1). Of the tested candidates for sacrificial layers, Micro-90 was chosen due to a straightforward lift-off process that can be accomplished within seconds by adding DI water. Parylene C deposition at a base pressure of 10 mTorr and deposition pressure of 25 mTorr resulted in conformal coating thickness across 6” wafers, and measured thicknesses were obtained by spectrophotometry and confirmed by profilometry. A combination of a top anti-reflective coating (TARC) and an optimized etching recipe was used to obtain a significantly lower pore expansion across samples, which were verified using scanning electron microscopy (**Figure S1 and S2**). Our results are analogous to previous works that included the use of hard masks that are costly, time-consuming, and require a chemically harsh removal step, leaving metallic residuals on the membranes and negatively impact subsequent use for cell culture. The lifted-off membranes are supported by SU-8 frames that provide large active areas (>1 cm2) and are easily transferable using tweezers (Figure 2A). SU-8 supports are used for easier handling and minimizing the deflection of membranes during transfer and use in cell culture.

**Figure 1.**
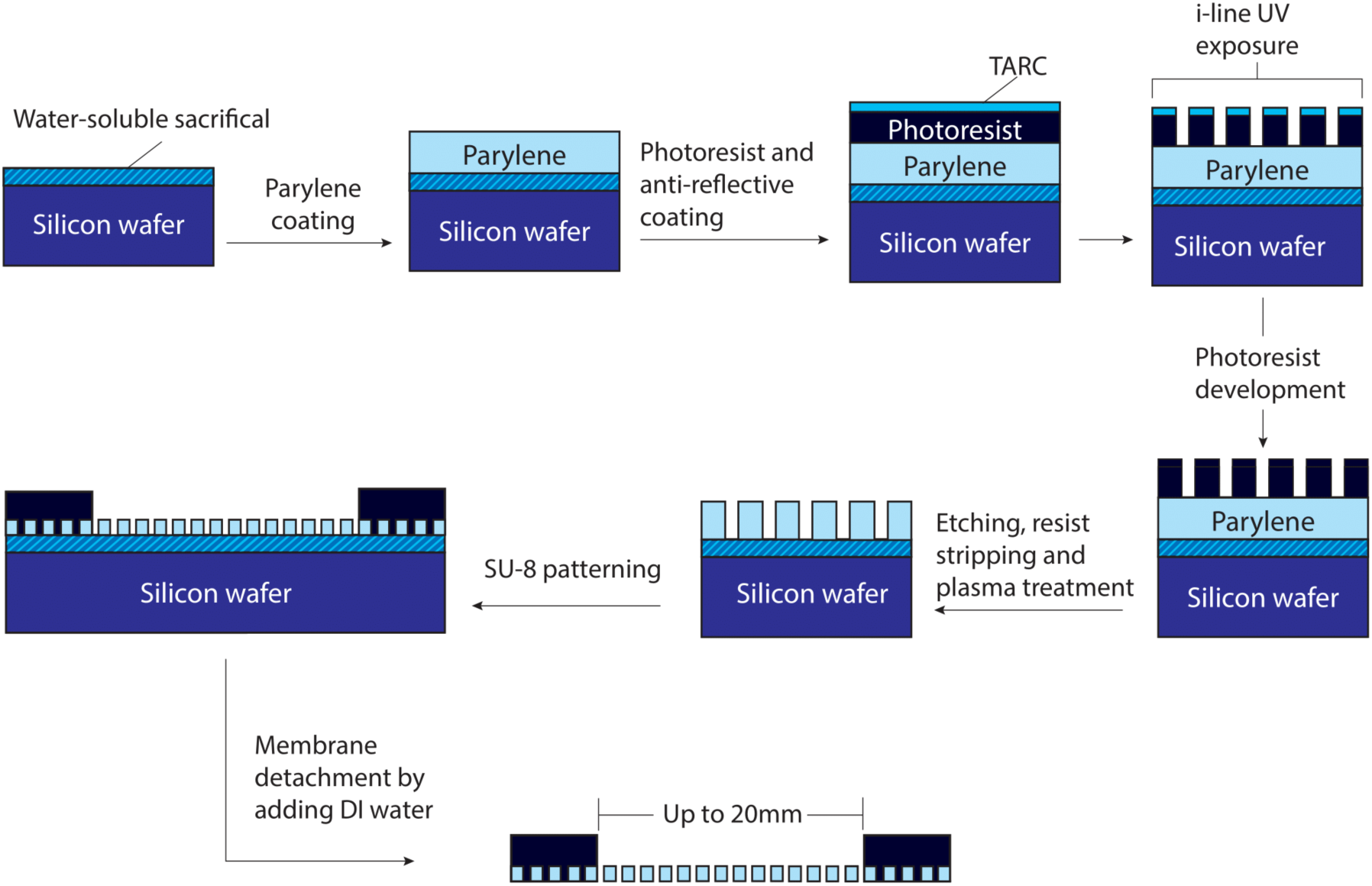
Microporous UPP membrane fabrication and introduction of a simple SU-8 frame to support robust handling and transfer. A step-by-step process for UPP membrane fabrication to achieve reliable pore geometries matching the mask design.

**Figure 2.**
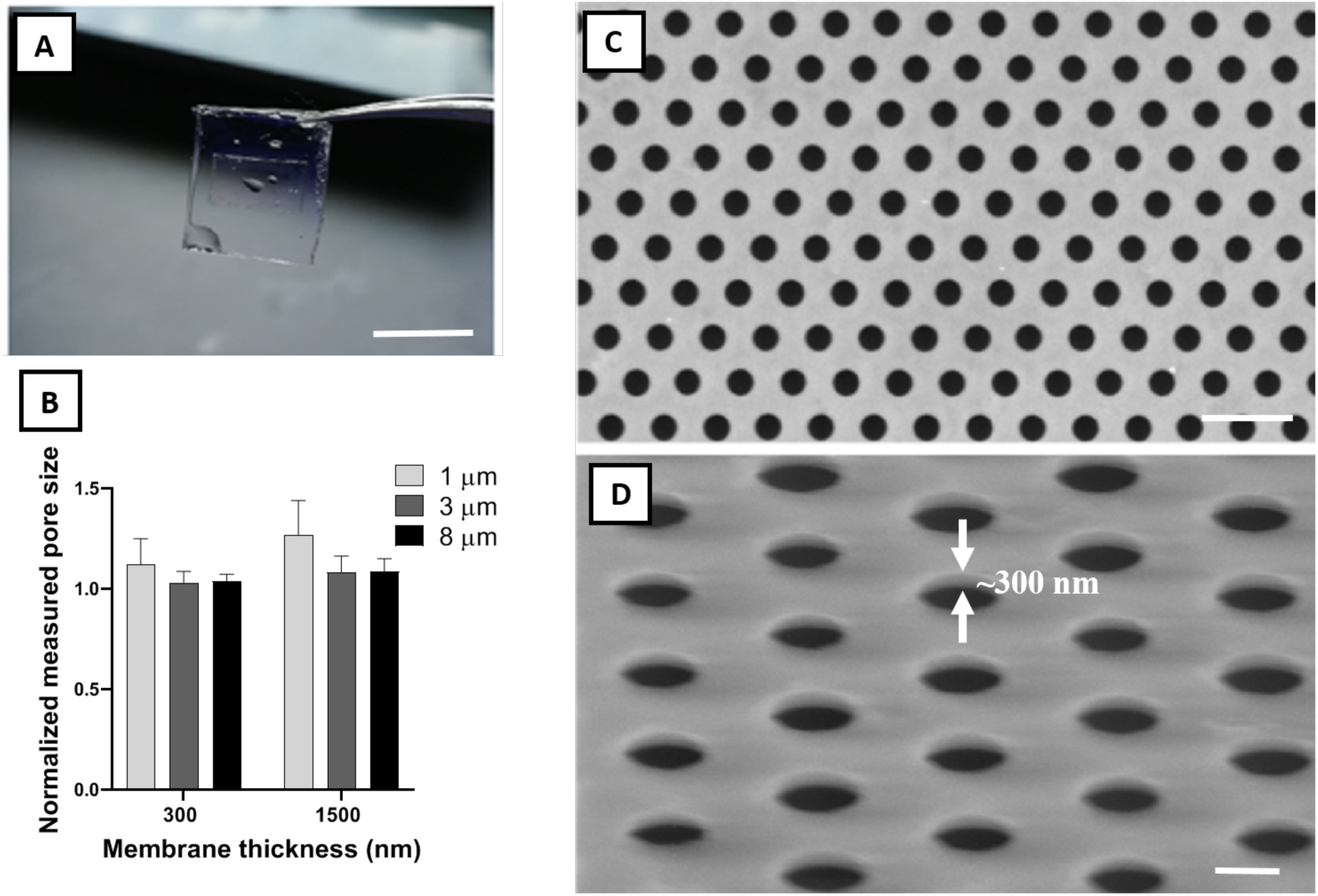
Physical characterization of frame-supported membranes. A) Water-facilitated release of a free-standing membrane with SU-8 support (scale bar=1 cm). B) Pore size fidelity (measured diameter divided by mask diameter) in fabricated membranes. Ideal pore size fidelity was generally observed in all samples. C) Top view and D) 60° tilted representative SEM images of a UPP membrane with 3 μm pore size and 300 nm thickness. White arrows demonstrate the ultrathin thickness of the membrane (~300 nm) (scale bar=10 & 4 μm, respectively).

Pore size fidelity relative to the mask design across three pore sizes (1, 3, and 8 µm) and two membrane thicknesses (300 nm and 1500 nm) revealed minimal pore size expansion under an optimized fabrication process (Figure 2B). Excellent pore size fidelity was observed in 300 and 1500 nm thick membranes with 3 and 8 µm pore sizes, and also in 300 nm thick parylene C membranes with 1 µm pore size. The 1500 nm thick parylene C membranes with 1 µm pore size showed an average pore expansion of 25%, which can be corrected by a change in mask design to account for the effect of higher thickness. SEM imaging shows uniform patterning across large areas with no noticeable defects (Figure 2C). The imaging also demonstrates the successful transfer of the pattern from the photoresist layer to the membrane layer with sharp edges (Figure 2D).

### 2.2. Thickness Gradient Membranes

A simple and inexpensive, custom-made polymethyl methacrylate (PMMA) chamber provides a reproducible system for creating a thickness gradient in the deposited membranes across the silicon wafer substrate, and the gradient can be easily tuned by changing the chamber width (Figure 3A). By modifying the opening distance between 1 and 4 mm, a thickness gradient of more than 200 nm/cm is achieved across a 2 cm vertical distance of the wafer (Figure 3B). An increase in the opening distance has a nearly linear correlation with the decrease in thickness gradient. A major challenge with fabricating porous membranes with a thickness gradient is that during the etching process the time required to create pores in the thicker regions will result in over-etching the thinner regions, leading to the expansion of the pores in the thinner regions. However, desirable pore size consistency was observed across the thickness gradient membrane, likely attributable to the combination of anisotropic etching and the use of TARC to minimize UV reflectance effect (Figure 3C).

**Figure 3.**
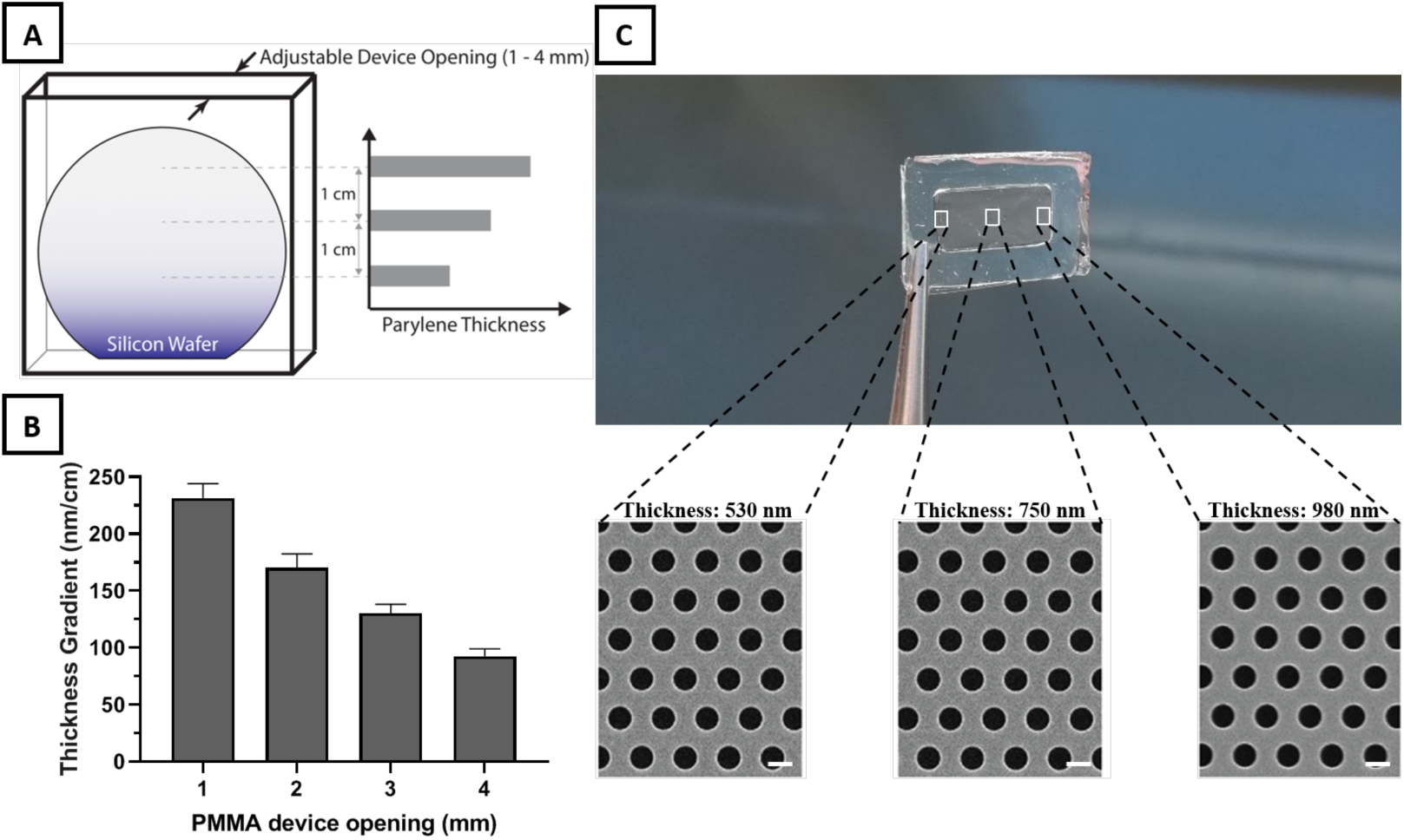
Thickness gradient membrane fabrication using a custom-made PMMA device. A) Illustration of the PMMA device designed with an adjustable top opening as the only path for parylene deposition on the enclosed silicon wafer. The restricted diffusion path for parylene monomers before polymerization at the wafer surface leads to a linear decrease in thickness from the top of the wafer down. B) Increases in the PMMA device opening results in a nearly linear decrease in thickness gradient across the membrane. C) Thickness difference across a frame-supported UPP membrane with a length of 2 cm. SEM images of different sites of frame supported membrane reveal a consistent pore size and distribution across different thicknesses (scale bar=1 μm)

### 2.3. Mechanical Testing

We next investigated the mechanical properties of UPP membranes at varying thickness and porosity. Previous studies that modeled mechanical properties of porous materials progressed from the simplistic assumption that no stress-concentration arises from pores (i.e. ultimate strength is correlated with volume fraction) to empirical models for material properties such as ultimate strength (σ_u_), such as the following equation:

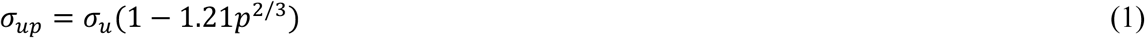

where *σ*_*up*_ and *p* are the ultimate strength and porosity of the porous material, respectively.^[39–42]^

Recent studies have performed various simulations and modeling and combined these to obtain a more accurate representation of material properties.^[43–47]^ They adopted the following equation:

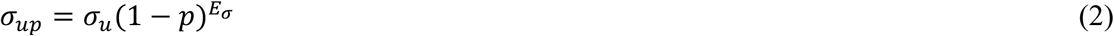

where *E*_*σ*_ is obtained through estimation and can be approximated as 2 based on theoretical and experimental data.^[44,45]^ Simplifying the equation yields:

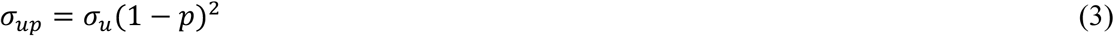

Although these models have been tested experimentally for characterizing the strength of a variety of materials with different shapes, such experimental validation has not been tested on ultrathin porous membranes. While there have been several notable studies that have investigated the mechanical characterization and tensile properties of non-porous thin films,^[48–51]^ we believe this is the first examination of the implications of introducing porosity in parylene nanomembranes. Previous studies on the mechanical properties of porous nanomembranes primarily used nanoindentation and bulge testing due to the challenge of conducting traditional tensile testing on ultrathin materials.^[52–54]^ To evaluate the mechanical properties of our UPP membranes, we applied a SU-8 grip support layer to both ends of membrane test strips, which enables evaluation of the membranes with a commercial tensile tester (**Figure S3**).

A summary of the results from the tensile tests is presented in Figure 4. Both the ultimate and yield strength follow a second-order dependence on the solid fraction of the membranes (1 − *p*), showing good agreement with the model described in Equation 3. Results for the yield strength are included here because this property is relevant to the membrane application of tissue culture. The intended purpose of UPP membranes is to act as a nominally two-dimensional substrate to enable facile imaging of cells using optical microscopy. If the membranes undergo significant plastic deformation from the stresses of handling and device integration and cell culture processes, then the membrane would no longer be flat and would diminish the ease of planar cell imaging.

**Figure 4.**
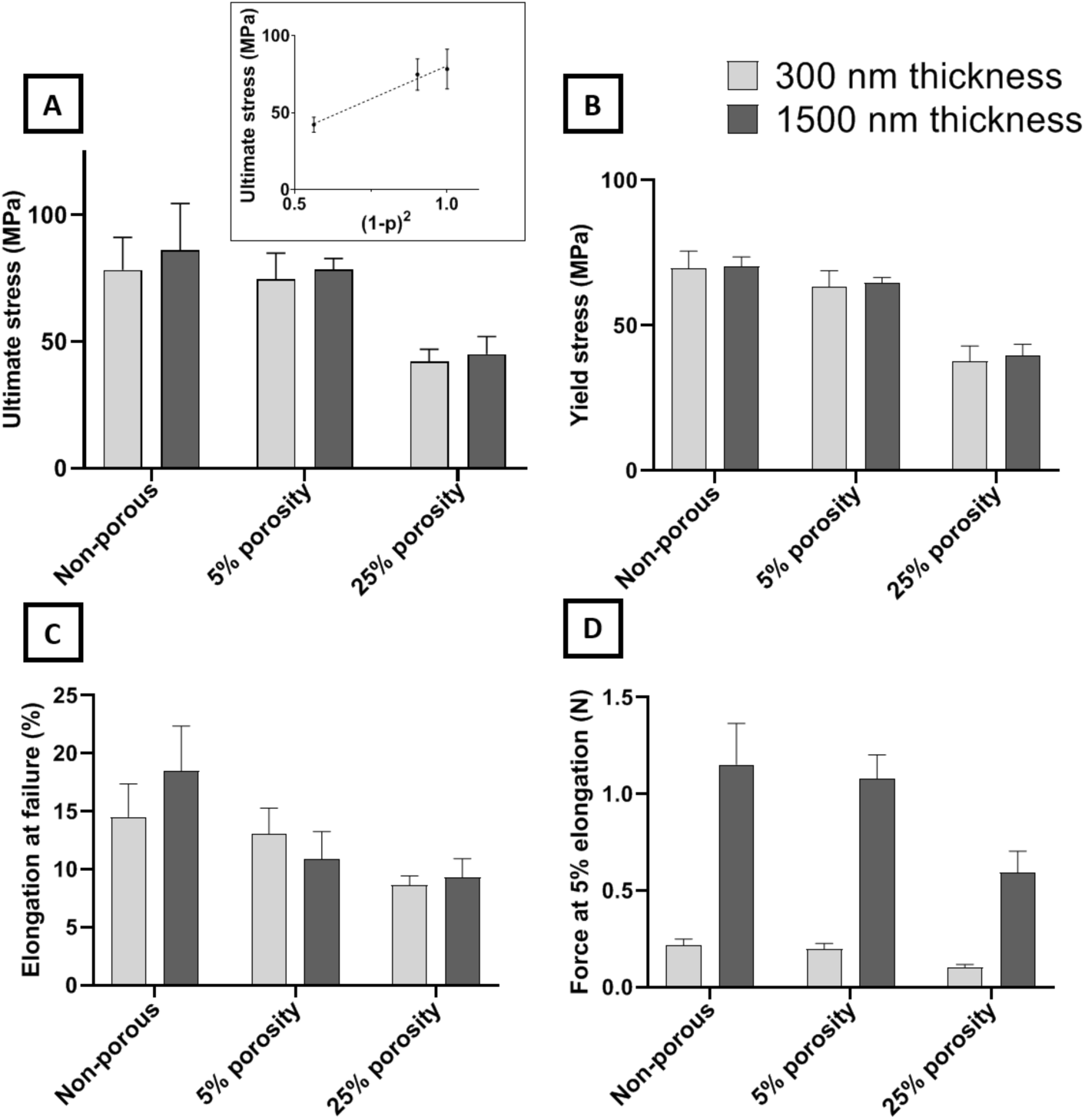
Mechanical characterization of ultrathin UPP membranes for various membrane designs (two thickness and three porosities). A) Ultimate stress of membranes. No significant difference was observed between thicknesses or between non-porous and 5% porosity. However, a ~47% decrease is observed by increasing the porosity to 25%. Ultimate stress is the plot as a function of Equation 3 in the inset. B) Yield stress of membranes. The trend of changes across different porosities was similar to the ultimate stress results. C) Elongation of the membrane at failure shows remarkable retention of resistance against mechanical strain after introducing 25% porosity while yielding no significant difference among different thicknesses. D) Force at a given elongation (5%) reveals the expected outcome that force is directly proportional to the thickness, and force decreases with increasing porosity, similarly to ultimate stress results.

Considering the apparent second-order dependence of the membrane strength on the solid fraction (i.e., (1 − *p*)), it is clear that the membranes should quickly become too weak at higher porosities. However, both the 5% and 25% porous membranes display a strength that is robust with respect to the stresses experienced in lift-off, mounting into tissue culture devices and cell culturing activities. Our experience even showed that membranes with porosities as high as 60% could be successfully lifted-off and transferred to cell culture devices, but subsequent handling to perform mechanical testing was not feasible due to their fragility. We targeted up to 25% porosity because it is more than double what is commonly available in track-etched membranes. Furthermore, we previously demonstrated that porous membranes with 25% porosity modulate cell-substrate interactions and cell behavior like softer, non-porous materials with Young’s moduli similar to *in vivo* tissue values.^[18]^

Theoretically, maximum elongation for all conditions should remain the same under ideal conditions. However, various defects can be introduced by increasing porosity and decreasing thickness that may not be accurately predicted nor avoided. This is even more pronounced when dealing with porous membranes of sub-micron thickness. Test results of the UPP membranes show that elongation at failure is essentially independent of the membrane thickness at all porosities. In terms of the effects of thickness on mechanical properties, no significant difference is observed between 300 nm and 1500 nm thicknesses for ultimate stress and elongation at failure. Additionally, the force required for 5% elongation is seen to scale directly with membrane thickness as expected.

In summary, mechanical characterization reveals that the strength characteristics of the UPP membranes follow an apparent second-order dependence on the solid fraction, which agrees with previous modeling and experimental work. The ultimate and yield strength exhibited by the 25% porous membrane is still at 50% of that of the non-porous one, and this is sufficient to ensure that the porous membrane is robust enough to handle the stresses of fabrication and cell culture.

### 2.4. Surface Roughness and Improved Cell Adhesion

Culture substrates are routinely modified to improve cell adhesion. Some approaches rely upon adsorption or attachment of extracellular matrix proteins or peptides, which can be expensive and have limited shelf-life.^[55,56]^ Alternatively, tissue culture polystyrene (TCPS) substrates are routinely treated with plasma.^[67]^ In order to improve cell adhesion to the UPP membranes, we explored treating the surface with oxygen plasma. We exposed membranes to two reactive ion etching (RIE) oxygen plasma powers (100 W and 200 W) and measured endothelial cell attachment after storing the membranes in atmospheric conditions for 1, 7, or 14 days. Both 100 W and 200 W resulted in significant improvements in cell adhesion over untreated membranes, increasing adhesion to levels similar to TCPS, which is consistent with previous studies involving surface modification of parylene substrates for improved cell adhesion (Figure 5).^[57,58]^ Another important aspect of these surface modifications is persistence over time. The advantages of the oxygen plasma treatment on some surfaces can be transient with hydrophobic recovery occurring within 24 hours.^[59]^ In our studies, we found treated UPP membranes maintained high levels of cell adhesion even after 14 days of storage in atmospheric conditions at room temperature.

**Figure 5.**
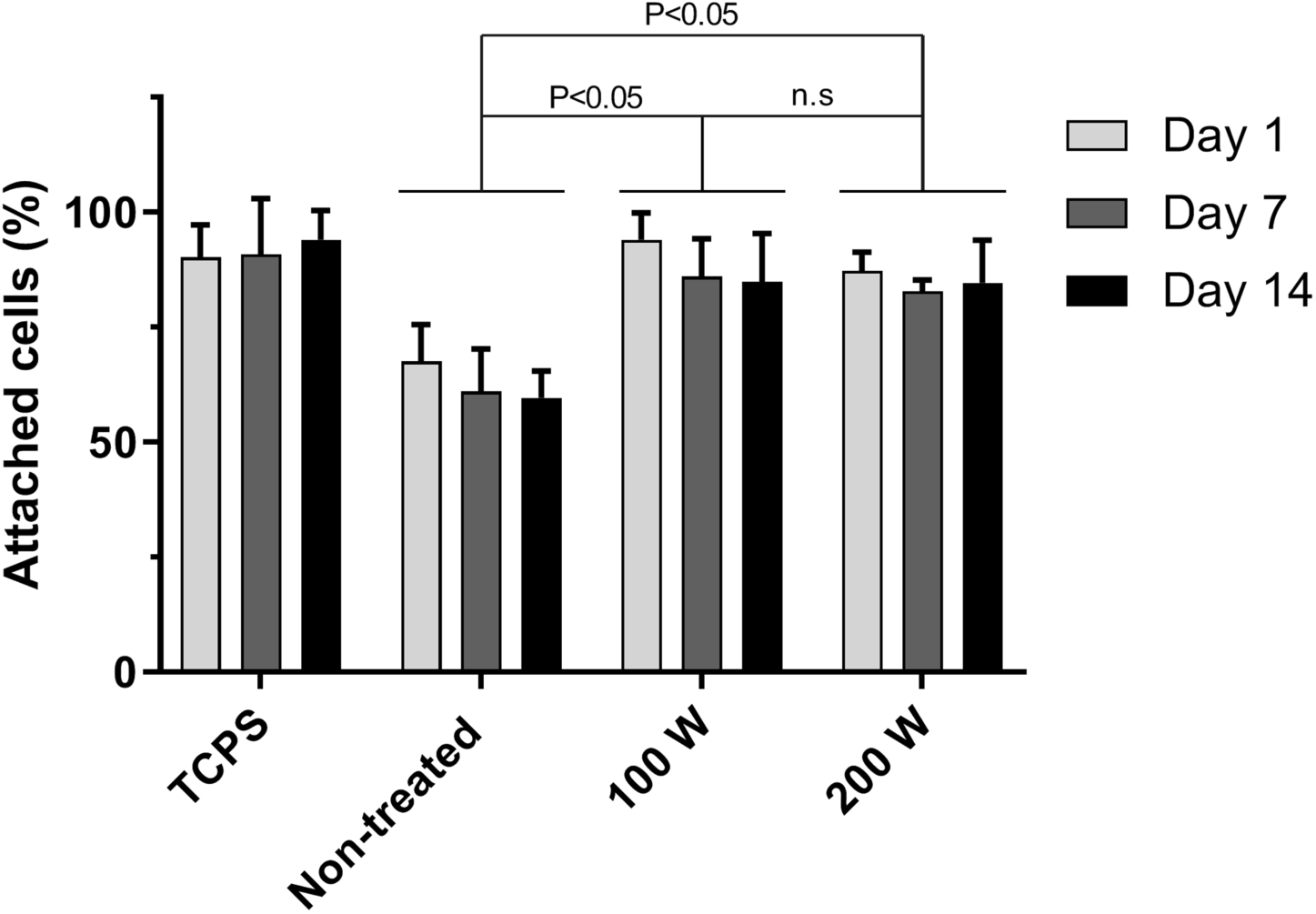
Modification of cell adhesion as a result of RIE plasma surface treatment. Membranes were treated with low power (100 W) and high power (200 W) RIE O2 plasma, and cell attachment was quantified for cells seeded on samples 1, 7, and 14 days after plasma treatment to assess any potential deterioration in the positive effect of plasma treatment over time. Use of RIE treatment led to a significant increase in the number of adhered cells as compared to non-treated membranes, leading to similar results of cell adhesion on tissue culture polystyrene, while the increasing RIE power from 100 W to 200 W led to no significant difference in cell adhesion. The positive effect of RIE treatments remained persistent until 14 days after the treatment.

Due to the persistent improvement in cell adhesion, we hypothesized that our plasma treatments may have induced a permanent physical change to the membrane surface. We used atomic force microscopy (AFM) to examine surface roughness of the UPP membranes before and after RIE plasma treatment. Our data show RIE treatment results in a rougher surface that may explain the improved and persistent cell adhesion (Figure 6). The changes are on a length scale that likely provides a greater surface area for protein adsorption, and may inhibit collagen motility, resulting in aggregation of collagen fibrils which can significantly increase cell gripping and adhesion as opposed to smooth collagen layer on smooth surfaces.^[57,60–62]^

**Figure 6.**
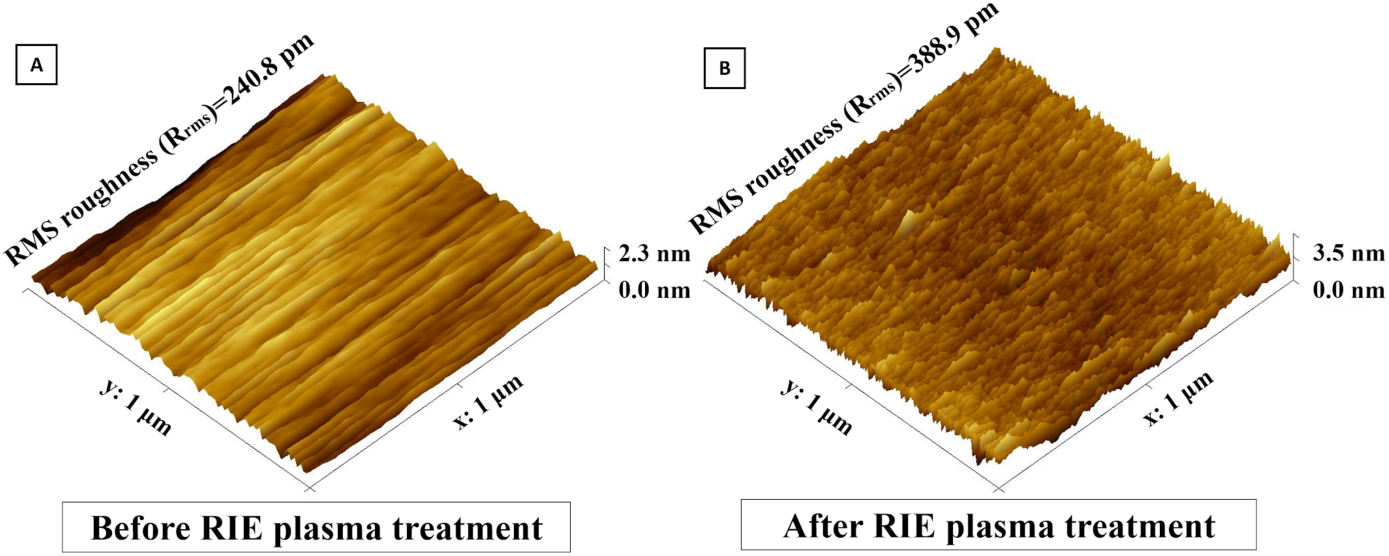
Surface analysis of UPP membranes before RIE plasma treatment and B) after RIE plasma treatment using AFM. Upon AFM analysis of the surface morphology, RMS roughness (R_rms_) was found to have significantly increased after plasma-treatment.

### 2.5. Potential applicability of UPPM to flow-based cell culture devices

Here, we aimed to show technical feasibility for UPP membrane integration into a prototype 3D-printed scaffold system for flow-based cell culture (Figure 7). To this end, frame-supported membranes were successfully seated on the top surface of the 3D-printed chamber (Figure 7B). Feasibility tests showed that the UPP membrane remained stable in place and did not tear when the flow was applied at a range of 1 to 20 ml/min using a pulsatile pump. When the pulsatile flow is applied, the circulating media did not diffuse underneath the UPP membrane.

**Figure 7.**
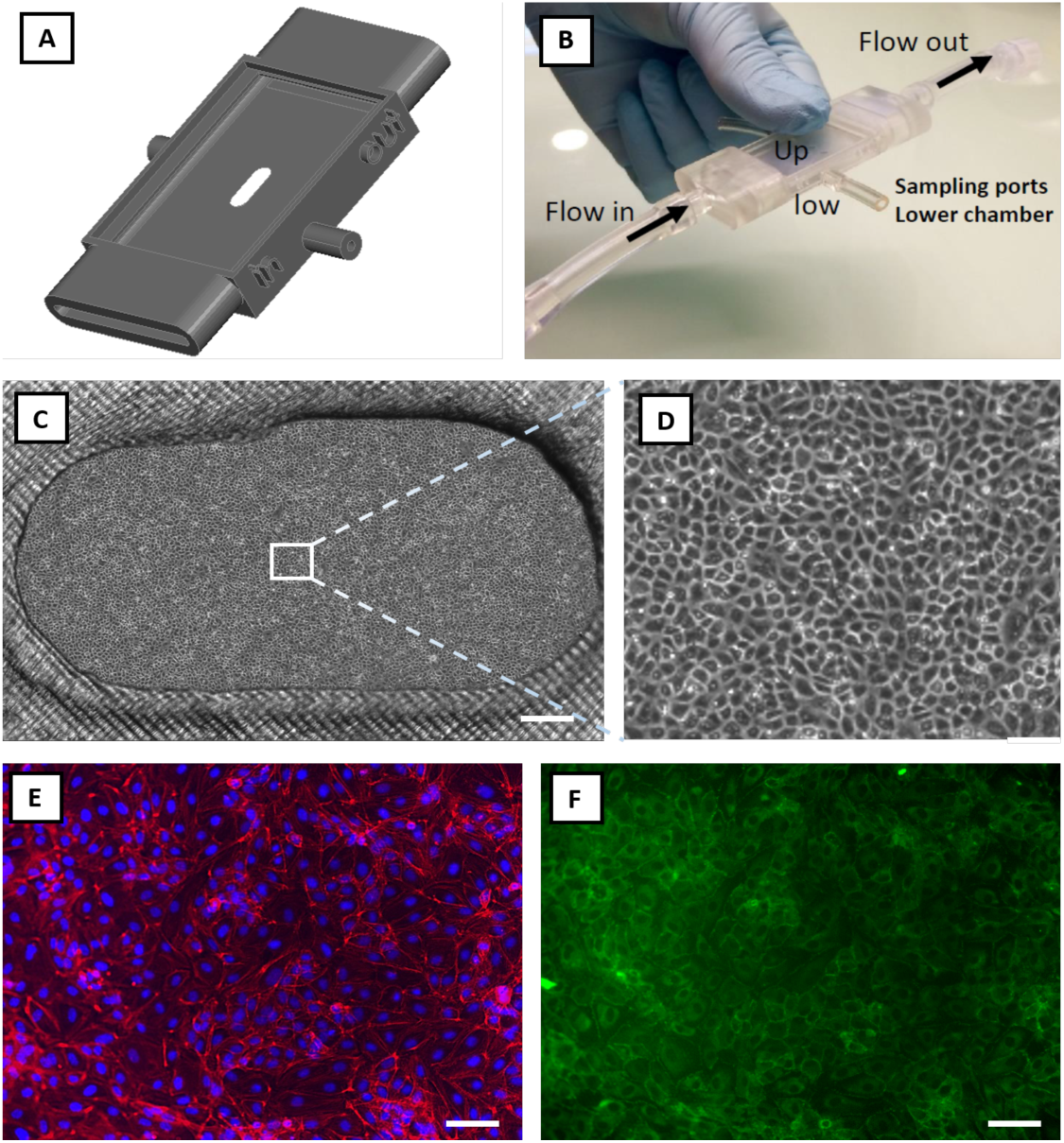
Feasibility study: integration of the UPP membranes into a prototyped flow-based *in vitro* model and cell culture viability. A) Graphical rendering of a 3D printed prototype connected to a peristaltic pump to provide pulsatile flow in the upper chamber (Up). B) The UPP membrane was successfully transferred on the upper chamber of the device, adhering and covering the whole surface, including the oval opening where the suspended membrane did not tear. The latter constitutes a proof-of-principle structural arrangement for co-culture development using UPP membranes. Under these conditions, the pulsatile flow did not detach or damage the UPP membrane, indicating the robustness of this system. C) Representative phase-contrast image of endothelial cells cultured on the UPP membrane in the prototype culture device (scale bar = 500 μm). D) Magnified optical image of the panel (C), showing confluent endothelial cells. E) Endothelial cells on UPP membranes visualized by DAPI (blue) and cytoskeleton (red) staining (scale bar = 50 μm). F) ZO-1 tight junction protein staining (scale bar = 50 μm).

Next, we tested whether a viable endothelial cell culture is attainable when UPP membranes are integrated into the 3D-printed chamber. Under these conditions, endothelial cells attached and grown to confluence within 3 days of seeding (Figure 7C-D). Phalloidin and nuclei DAPI staining showed physiological cytoskeletal organization and cell morphology on the UPP membranes (Figure 7E). Tight junction protein staining (Figure 7F) indicates the development of continuous endothelial intercellular contacts supporting the potential applicability of UPP membranes to cell transmigration or drug transport studies. Taken together, these results advocate for the full development and testing of UPP membrane-based *in vitro* culture or co-culture devices, including applicability to flow-based systems.

## 3. Conclusion

In summary, we developed methods to efficiently fabricate robust UPP membranes that can be used for viable cell culture and integrated into prototyped *in vitro* devices. UPP membranes can be produced with variable and even gradient thicknesses to investigate cell-cell interactions. Our study on membrane thickness, porosity, and mechanical properties demonstrated agreement with the existing empirical models used to predict practical upper limits of porosity and lower limits of thickness. Lastly, we showed that oxygen plasma etching improves cell adhesion on UPP membranes. Further studies are now needed to prove the biological advantages and the *in vivo* mimicry associated with the integration of UPP membranes into *in vitro* devices or chips.

## 4. Experimental Section

### Sacrificial layer deposition and parylene coating

Micro-90 was used as a water-soluble sacrificial layer. This was achieved by spin-coating the solution on top of a 6-inch (100) silicon wafer (500 rpm for 3 s, followed by 3000 rpm for 45 s). The last step in the fabrication process requires simply exposing the attached membrane to Deionized (DI) Water, which dissolves Micro-90. A solution of Micro-90 diluted to 5% in DI water was identified as the optimal concentration. Although higher concentrations of Micro-90 led to much easier detachment, they often resulted in delamination during the fabrication process.

Parylene-C coating was done using DPX-C dimer (Specialty Coating Systems, USA) in an SCS Labcoter® 2 parylene deposition system (PDS 2010, Specialty Coating Systems, USA). Briefly, silicon wafers are placed in the deposition chamber with an optimized separation distance (minimum of 10 cm) to ensure thickness uniformity across the wafer, and dimer is loaded in the vaporizer (nominally 800 nm final parylene C thickness per 1 gram of dimer). The process begins at a base chamber pressure of 10 mTorr, and the dimer-cracking furnace is heated to 690 °C. Then, the vaporizer is ramped to a final temperature of 175 °C, causing the sublimation of dimer. The temperature ramp rate of the vaporizer is controlled to maintain a chamber pressure of 25 mTorr. Thickness measurements were performed by both Tencor P2 long scan profilometer (KLA-Tencor, USA) and NanoSpec Spectrophotometer (Nanometrics Incorporated, USA). Membranes with a thickness gradient across the diameter of the wafer were obtained by placing the wafer in a custom-made polymethyl methacrylate (PMMA) fixture (Figure 3A) for the deposition. The PMMA fixture creates a narrow slot that restricts the diffusion of the parylene monomers and thereby reduces the likelihood of the monomers reaching the wafer surface toward the bottom of the fixture. The gap between the wafer and the fixture wall can be adjusted allowing for a range of thickness variations.

### I-line photolithography

HDMS prime (Microchemicals, Germany) was used as an adhesion promoter between parylene and photoresist, and was applied by spin-coating at 3000 rpm for 45 s followed by baking at 140 °C for 1 min. AZ MiR 701 positive photoresist (14 cPs) (Microchemicals, Germany) was applied by spin-coating at 1500 rpm for 45 s followed by soft-baking at 90 °C for 1 min to obtain a thickness of 1.5 µm. To prevent UV light reflection which leads to undesired pore size change, AZ Aquatar (Microchemicals, Germany) was used as a top anti-reflective coating (TARC) with a manufacturer-recommended spin-coating recipe of 500 rpm for 3 s followed by 2200 rpm for 30 s. The advantage of using TARC over bottom anti-reflective coating (BARC) such as AZ Barli II is the removal during the developing step without the necessity of separate etching steps while obtaining comparable results.

Photoresist exposure was performed on an ASML PAS 5000 i-line 5X Stepper (ASML, Netherlands). In order to save time and efforts and minimize associated costs, multiple designs with different final pore sizes and pore spacings (1 µm, 1 µm, 3 µm, and 8 µm pore size with 2 µm, 8 µm, 6 µm, and 16 µm center to center spacings, respectively) were embedded in a single mask. Multiple parameters including exposure dose and focus offset were adjusted and tested for each patterned image. The wafers then went through a post-exposure bake at 110 °C for 1 min. The exposed photoresist was developed using Microposit MF CD-26 developer (Microchemicals, Germany) followed by hard-baking at 140 °C for 1 min. All photolithography steps were performed while the ambient cleanroom humidity was within a 38-42% range.

### Reactive ion etching

Previous works involving etching parylene mainly used Reactive Ion Etching (RIE) with Oxygen to etch parylene. However, creating high-resolution patterns with oxygen etching proved to be challenging due to a generally higher etching rate of the photoresist versus the parylene and isotropic etching of both the parylene and photoresist layers. A variety of etching recipes were evaluated using a Trion Phantom 3 system (Trion Technology, Inc., USA), and a combination of 50 sccm of Oxygen (O2), 5 sccm of Sulfur hexafluoride (SF6), and 5 sccm of Argon (Ar) proved to be an optimal etching recipe (Figure S1). The chamber pressure was kept at 50 mTorr and power was set at 175 W with a tolerance of 2 mTorr and 5 W, respectively. The residual photoresist was removed by immersing that wafer in the PRS-2000 solution (JT Baker, Inc. USA) at 75 °C for 10 min.

### Membrane frame design and embedment

To enable easier transfer of the UPP membranes after lift-off, a “scaffold structure” was patterned on top of the UPP membrane through contact lithography of SU-8 to obtain frames with different designs. Wafers were dehydrated at 150 °C for 10 min. Then, SU-8 3025 was deposited with a spin-coating recipe of 500 rpm for 10 s followed by 3000 rpm for 45 s, soft-baked at 95 °C for 10 min, and transferred to a cool plate, which resulted in a SU-8 layer with 20 µm thickness. The wafers were exposed in contact lithography to broadband spectrum UV light for exposure of 250 mJ/cm2. Post-exposure bake consisted of 65 °C for 1 min, cool plate for 10 min, and 95 °C for 5 min. Wafers were developed using SU-8 developer (Microchemicals, Germany) for 5 min with mild agitation. The wafers were then rinsed with IPA for 10 sec, dried, and given a final hard bake at 120 °C for 150 s.

### Membrane lift-off and characterization

The wafers were cleaved into nominal 2 cm by 3 cm samples before membrane lift-off. The cleaved wafers were simply soaked in DI water for 1-2 minutes which dissolved the sacrificial layer and facilitated membrane lift-off. The frame-supported membranes were lifted with tweezers to transfer to other containers for subsequent characterization or incorporation into cell culture devices. The physical properties of the membranes were characterized by scanning electron microscopy (SEM) analysis.

### Mechanical characterization

In order to quantify the impact of porosity, pore size, and thickness on the mechanical properties of the porous membranes, 1 cm ⨉ 3 cm rectangles of UPP membranes were fabricated with thick SU-8 support regions covering 5-mm strips on either end to serve as gripping area supports. A CellScale® UniVert Biomaterial Tester (CellScale, Canada) equipped with a 10 N load cell was used for mechanical tensile testing. The SU-8 regions were held between Polyoxymethylene (POM) clamps specifically designed for soft materials. CellScale analysis software was used to extract ultimate stress, elongation at failure, and force at 5% elongation for three different porosities (0%, 5%, and 25% porosities) and two thicknesses (300 nm and 1500 nm). A minimum of five samples was used for each condition.

### Cell adhesion

Human Umbilical Vein Endothelial Cells (HUVECs) (Thermo Fisher Scientific, USA) used for biocompatibility and cell attachment were passages 4-6. They were grown in M200 media, supplemented with 2% Large Vessel Endothelial Supplement (LVES), 100 μg/ml penicillin, and 100 μg/ml streptomycin.

RIE oxygen plasma treatment was used to modify the surface for better cell attachment. After completion of fabrication steps and before lift-off, membranes were exposed to RIE oxygen plasma in the Trion phantom III system (40 sccm, 50 mTorr) with 100 and 200 W power. Membranes were washed in 70% ethanol for 30 min and subsequently in ultrapure water for 30 min, all done in a laminar-flow biosafety cabinet after lift-off. Cells were seeded on the membranes, and tissue culture polystyrene (TCPS) as control with a density of 5000 cells/cm2. TCPS served as a typical standard cell culture substrate with adequate cell adhesion. The total number of attached cells was counted on membrane and control samples were counted after 6 hr. Although oxygen plasma treatment has been previously used for cell culture membranes for enhancing cell attachment, its effects and stability over time have not been quantified. To quantify its effects on UPP membranes, cell seeding was performed after 1, 7, and 14 days after oxygen plasma.

### Device fabrication and assembly

Cell culture devices were designed using SolidWorks® software and printed using Form 2 Stereolithography (SLA) printer (Formlabs Inc., USA) with a layer precision of 25 μm. This was followed by thorough washing with isopropyl alcohol (IPA) and curing at 60 ℃ for 45 min. After membrane lift-off using DI water, they were transferred using a tweezer to the 3D printed device. PBS was added on top of the membrane and left to dry to promote adhesion between the membrane and the device.

### Cell culture and immunofluorescence

Bovine aortic endothelial cells (BAECs) were cultured in Dulbecco’s Modified Eagle Media (DMEM) supplemented with 10% FBS, 1% L-glutamine, 100 μg/ml penicillin, and 100 μg/ml streptomycin. Cells were detached and sub-cultured per manufacturer’s instructions using TrypLE. BAEC media was exchanged every 2-3 days and cells were passaged at 80% confluence. BAECs were used between passages 3-4. Cells were seeded on the samples with a seeding density of 10,000 cells/cm2 in a 200 μl cell solution and left for 4 hr before flooding with media. After 3 days, cells were fixed with 3.7% formaldehyde for 15 minutes, washed three times with PBS, and then permeabilized with 0.1% Triton X-100 for 3 minutes. Cells were blocked with 2% BSA for 15 minutes and again washed with PBS. For visualization of nuclei and stress fibers, cells were stained with DAPI (300 nM), and 1:400 AlexaFluor 488 conjugated phalloidin. For visualization of tight junctional protein ZO-1, the cells were stained with 1:100 AlexaFluor488 conjugated anti-ZO-1/TJP1, Clone ZO1-1A12 (Affymetrix eBioscience, San Diego, CA).

## Acknowledgments

The authors acknowledge support by NIGMS of the National Institutes of Health (R35GM119623) to T.R.G. We would like to thank Julia Lunavictoria for help with illustrations. We thank Sean O’Brien, Patricia Meller, and John Nash from Semiconductor and Microsystems Fabrication Laboratory (SMFL) at Rochester Institute of Technology, and James A. Mitchell and Nursah Kokbudak from URnano cleanroom facilities at the University of Rochester. We would like to thank Frederic deBock and Aphrodite Alverti-Vallerand (Institute of Functional Genomics, France) for technical assistance with tissue culture and the initial set-up of 3D printing. The content of this publication is solely the responsibility of the authors and does not necessarily represent the official views of the National Institutes of Health.

