## Supplemental Figures for "Robust and Gradient Thickness Porous Membranes for *In Vitro* Modeling of Physiological Barriers"

Supporting Information: 3 Figures.

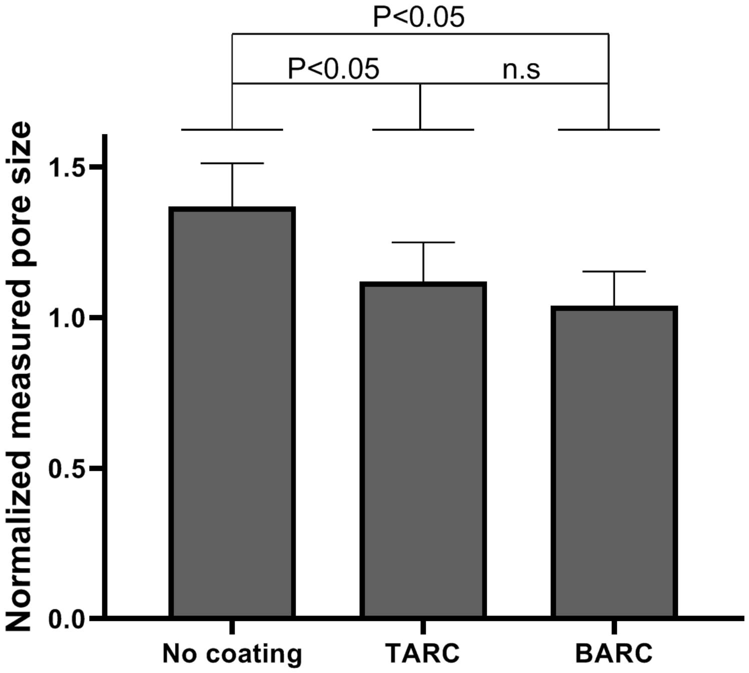


Figure S1. Effect of anti-reflective coating on ultrathin porous parylene C (UPP) membrane pore expansion after patterning and etching. The use of TARC on the photoresist layer resulted in significantly lower pore size (indicative of size fidelity to the mask design), while there was no significant difference between applying TARC and BARC. Considering the simplicity of using TARC, this is highly desirable.

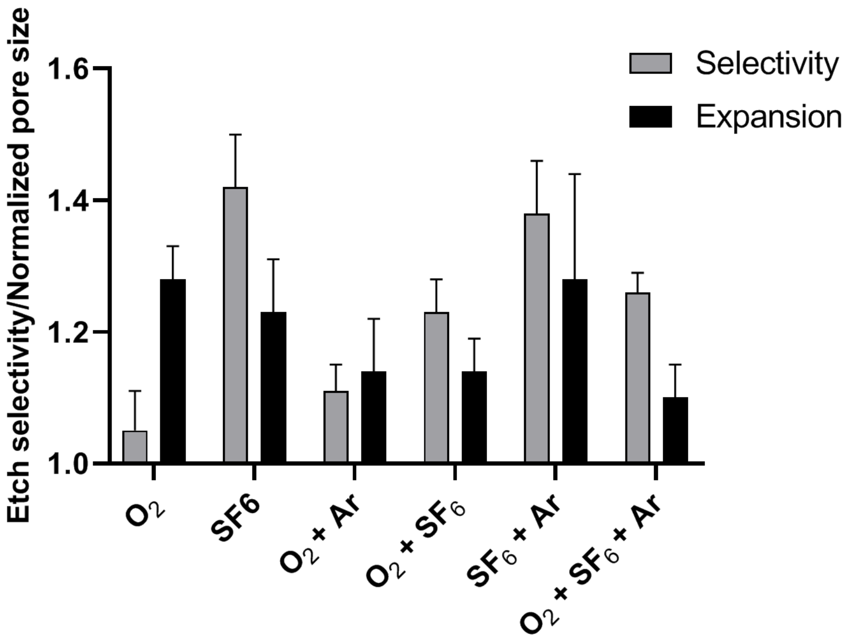


Figure S2. Parylene to photoresist etching selectivity and pore size expansion under different etching recipes. Different combinations of $O_2$ (50 sccm), Ar (5 sccm), and $SF_6$ (5 sccm) were tested to obtain the highest selectivity of parylene etching rate divided by photoresist etching rate and lowest pore expansion characterized by SEM. Although $SF_6$ and $SF_6$ + Ar led to the highest selectivity, they led to undesirable pore expansion. $O_2$ + $SF_6$ + Ar resulted in the best pore expansion performance while yielding fairly good selectivity.

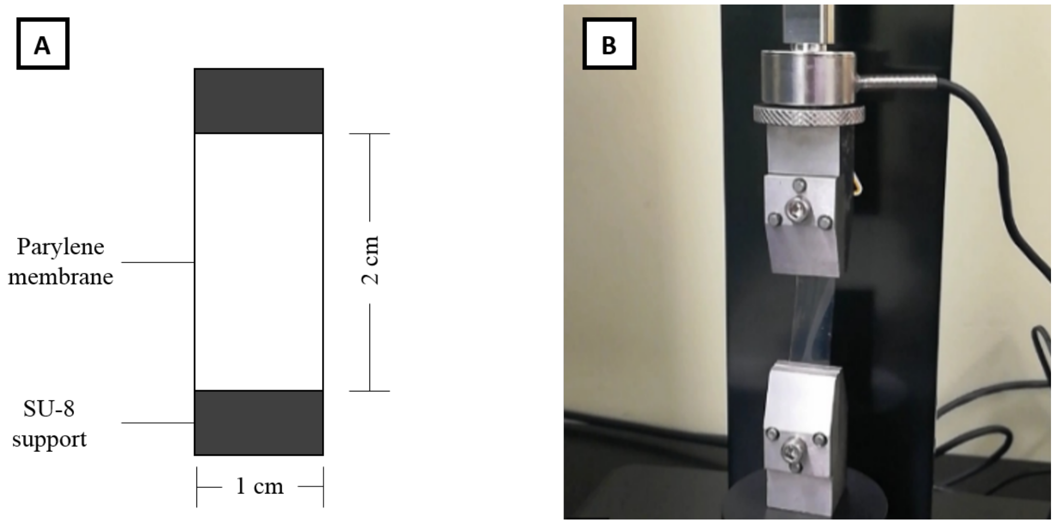


Figure S3. Mechanical characterization setup. A) Blocks of SU-8 were patterned on the UPP membrane before membrane lift-off. B) Upon membrane lift-off, SU-8 supported membrane is carefully transferred to the tool with tweezers and SU-8 supports were used as gripping regions in the CellScale® UniVert Biomaterial Tester.
